# ReLo is a simple and quick colocalization assay to identify and characterize direct protein-protein interactions

**DOI:** 10.1101/2022.03.04.482790

**Authors:** Harpreet Kaur Salgania, Jutta Metz, Mandy Jeske

## Abstract

The characterization of protein-protein interactions (PPIs) is fundamental for understanding biochemical processes. Many methods have been established to identify and study direct PPIs; however, the screening and investigation of PPIs involving large or poorly soluble proteins remain challenging. As a result, we developed ReLo, a simple, rapid, and versatile cell culture-based method for detecting and investigating interactions in a cellular context. Importantly, our data strongly suggest that with ReLo specifically direct binary PPIs are detected. By applying additional bridging experiments ReLo can also be used to determine the binding topology of subunits within multiprotein complexes. Moreover, ReLo has the potential to identify protein domains that mediate complex formation, screen for interfering point mutations, study interactions that depend on conformation or protein arginine methylation, and it is sensitive to drugs that mediate or interfere with an interaction. Taken together, ReLo is a simple and quick alternative for the study of PPIs particularly when established methods fail.

## INTRODUCTION

The identification and characterization of protein-protein interactions (PPIs) are a routine laboratory practice and lay the foundation for understanding biological processes. PPIs can be identified through various established, mass spectrometry-coupled screening methods, including coimmunoprecipitation (co-IP), tandem affinity purification, and proximity-dependent labeling approaches, such as BioID or APEX ^1–6^. Thus, a list of interacting protein candidates is obtained, and these candidates are usually ranked according to their abundance in eluate fractions. Determining which of these candidates are truly direct binding partners requires subsequent validation experiments, often involving *in vitro* methods, such as GST pull-down assays, which depend upon the availability of purified proteins. If the proteins are poorly soluble and cannot be obtained through recombinant protein expression or if expertise in recombinant protein expression and purification methods is lacking, PPIs can be validated using cell-based assays.

Yeast two-hybrid (Y2H) and protein complementation assays (PCA) are well-established techniques in which an interaction results in the reconstitution and subsequent detection of a split reporter protein, such as a transcription factor, ubiquitin, an enzyme, or a fluorescent protein ^7–12^. However, standard Y2H and PCA assays are not particularly suitable for the analysis of potentially unstable proteins, as they are barely expressed or rapidly degraded in a cell, resulting in unreliable, mostly false-negative results ^7, 10^. Therefore, to obtain conclusive findings, negative results require additional assessment of the protein expression levels, which complicates the process, particularly when probing many PPIs.

Cell-based PPI methods that are better suited for testing interactions involving potentially unstable proteins are based on fluorescent protein tagging and colocalization readouts and thus allow the simultaneous monitoring of both PPIs and protein expression levels via fluorescence microscopy. The readout of these colocalization assays is usually the translocation of one protein upon its association with a second distinctly localized protein (e.g., localization to a membrane, the nucleus, or granules). ’Cytoskeleton-based assay for protein-protein interaction’ (CAPPI), ’membrane recruitment assay’ (MeRA), and ’knocksideways in plants’ (KSP) are translocation assays developed for use with plant cells ^13–15^. ’Nuclear translocation assay’ (NTA), ’emerging circle of interactive proteins at specific endosomes’ (ECLIPSE), and ’protein interactions from imaging of complexes after translocation’ (PICT) are assays that require the addition of a compound (e.g., rapamycin) to monitor the translocation upon a PPI ^16–18^. Other translocation assays are based on oligomerization/aggregation readouts ^19–21^ and may thus not be suitable for studying interactions with proteins that form granules on their own within a cell. Importantly, none of the described translocation assays have been assessed based on their ability to distinguish direct binary interaction from those where the two proteins tested are eventually bridged by cell-endogenous proteins.

We developed a simple and fast translocation PPI assay named ReLo for use with an animal cell culture. This assay is based on the relocalization of a protein upon its interaction with a second membrane-anchored protein. We applied ReLo to many large proteins, most of which harbor long disordered regions and are known to be insoluble after recombinant protein expression experiments. Using this set of proteins, we demonstrate that ReLo can be employed to identify and characterize PPIs. Importantly, using two structurally well-characterized multidomain protein complexes, we show that only direct interactions are detected using ReLo, a prerequisite for the analysis of previously unknown protein complexes through *in vitro* and structural biology methods. Furthermore, ReLo is responsive to drug treatment, which enables the study of drug-induced interactions and the screening of small PPI inhibitors. In summary, ReLo is a simple, rapid and versatile tool that enables a thorough initial description of direct PPI networks.

## RESULTS

### ReLo: a simple and robust cell culture-based PPI assay

As a preparation for the ReLo assay, two proteins of interest were fused to a red fluorescent protein (mCherry) and a green fluorescent protein (EGPF, mEGFP). Importantly, one of the constructs carried an additional fusion to a membrane-anchoring protein domain, which led to distinct subcellular membrane localization of the fusion protein. Upon interaction, the second protein is expected to colocalize with the anchored protein to the membrane and it would thereby relocalize with respect to its original location (**Fig. 1a)**. Thus, we refer to the assay as the ’relocalization PPI assay’, abbreviated ’ReLo’.

**Fig. 1.**
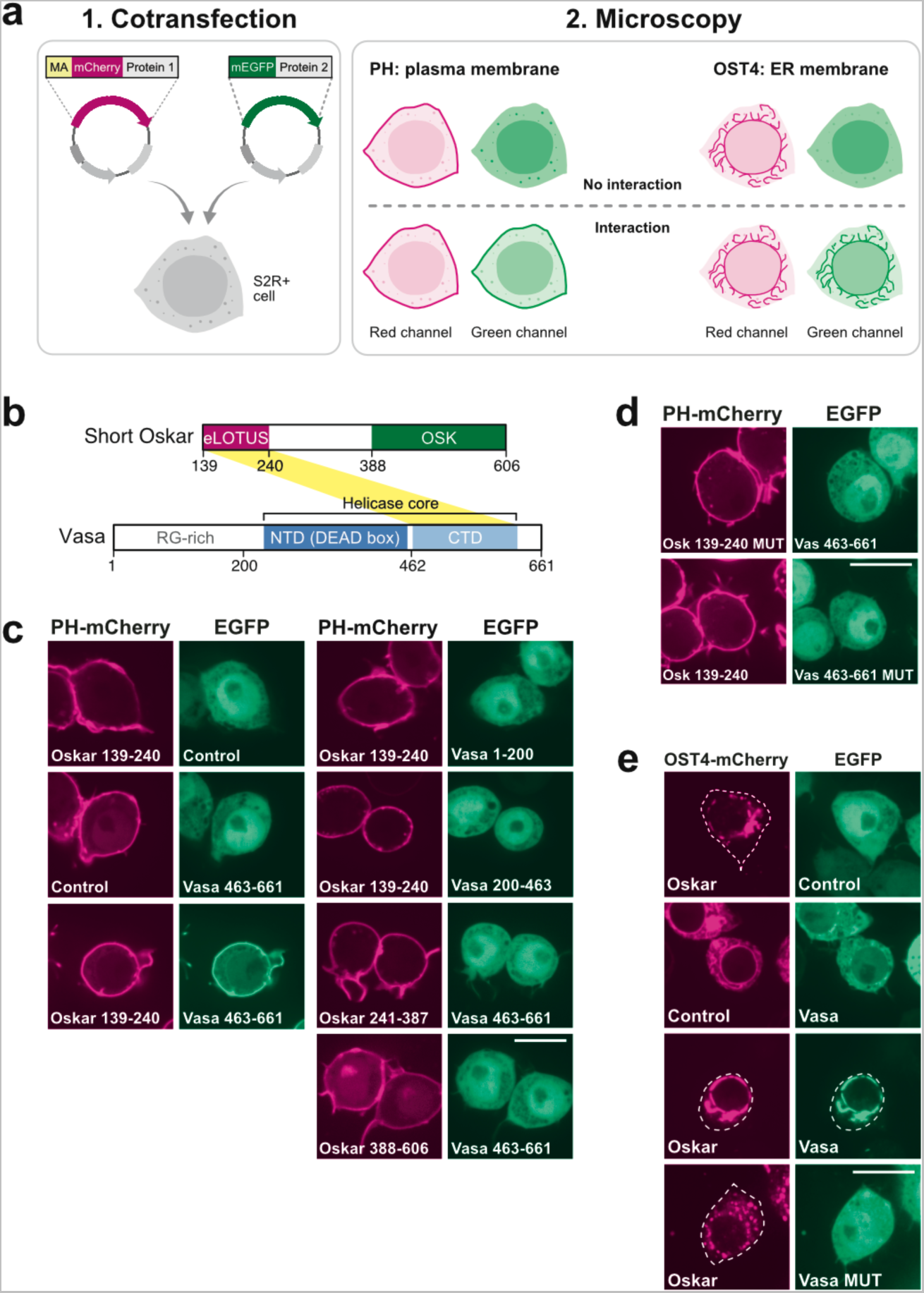
ReLo assay and its use in PPI mapping and mutational analysis. **a** Plasmids encoding fluorescently tagged proteins 1 and 2 are cotransfected into S2R+ cells. Protein 1 carries an additional fusion to a membrane anchoring (MA) domain, i.e., PH or OST4. After 48 h, protein localization is analyzed by confocal fluorescence microscopy. If protein 2 interacts with protein 1, protein 2 is relocalized to the plasma membrane (PH domain) or ER (OST4). **b** Domain organization of *Drosophila* Oskar and Vasa proteins. Oskar is expressed in two isoforms, and Short Oskar lacks amino acids 1-138. The eLOTUS domain of Oskar was previously shown to interact with the C-terminal domain (CTD) of Vasa (yellow stripe), and the crystal structure of the complex has been resolved ^28, 30^. **c** Vasa 463-661, but not Vasa 1-200 or 200-463, interacted with the Oskar eLOTUS domain. Vasa 463-661 did not interact with Oskar 241-396 or Oskar 398-606. **d** Oskar A162E/L228E and Vasa F504E point mutations (MUT) interfered with the Oskar-Vasa interaction. **e** OST4-mCherry Oskar localized to the ER (top panel), and Vasa fully relocalized with Oskar to the ER (bottom panel), while Vasa MUT did not. In the “Control” experiments the respective construct was coexpressed with the vector indicated at the top of the panel lacking an insert. The scale bar is 10 µm.

ReLo is based on a simple methodology in which cells are seeded onto 4-well chambered coverslips and cotransfected with the desired combination of plasmids. After 48 h, the protein localization is analyzed by live-cell confocal fluorescence microscopy (**Fig. 1a)**. We used S2R+ cells, which were derived from semiadherent Schneider’s-line-2 (S2) cells, which in turn have been established from late *Drosophila* embryos ^22, 23^. Compared with S2 cells, S2R+ cells show greater adherence to dishes, and thus they do not require coated dishes for their adhesion prior to live-cell microscopy. All ReLo plasmids carry an in-frame blunt end restriction site, allowing the application of a simple, fast, and straightforward cloning procedure (see Methods section).

For the anchoring of cytoplasmic proteins of interest to a membrane, we selected the pleckstrin homology (PH) domain of rat phospholipase Cο_1_ (PLCο_1_), which specifically recognizes phosphatidylinositol 4,5-bisphosphate ^24, 25^ and thus directs the fusion construct to the plasma membrane of a cell (**Fig. 1a)**. In the ReLo assay, the membrane localization was independent of whether the PH domain was fused to the N- or C-terminus of a protein (**Supplementary Fig. 1a)**. Unfortunately, nuclear proteins fused to the PH domain were only inefficiently retained within the cytoplasm and barely localized to the plasma membrane (**Supplementary Fig. 1c)**. Hence, the testing of PPIs to a nuclear protein may result in false-negative outcomes if the interaction partner resides in the cytoplasm. Therefore, we tested alternative membrane-anchoring domains to assess their ability to retain nuclear proteins within the cytoplasm. Hence, we selected the minimembrane protein subunit 4 of the yeast oligosaccharyltransferase complex (OST4), which was previously used as N-terminal fusion protein to localize a nuclear protein to the endoplasmic reticulum (ER) ^26, 27^ (**Supplementary Fig. 1d**). Because the plasma membrane localization of the PH domain distinguishes from a ubiquitous cytoplasmic localization better than the ER localization of OST4, we prefer using the PH domain in ReLo whenever possible.

### PPI mapping and mutational analysis

As a proof of concept for interaction studies with ReLo, we tested a previously characterized protein complex consisting of the extended LOTUS (eLOTUS) domain of Oskar (Oskar 139-240) and the C-terminal RecA-like domain (CTD) of the ATP-dependent DEAD-box RNA helicase Vasa (Vasa-CTD, Vasa 463-661) ^28^ (**Fig. 1b)**. Only the short isoform of Oskar (Short Oskar, aa 139-606) interacts with Vasa ^29^, and therefore, only Short Oskar and its domains were used in the following experiments (**Fig. 1b)**. In our setup, the eLOTUS domain of Oskar was fused to PH-mCherry and localized to the plasma membrane. Vasa-CTD was fused to EGFP and localized ubiquitously within the cytoplasm and nucleus. When coexpressed with PH-mCherry-eLOTUS but not with PH-mCherry alone, EGFP-Vasa-CTD relocalized to the plasma membrane (**Fig. 1c)**. The unstructured region of Vasa (Vasa 1-200) and the N-terminal RecA-like domain (Vasa 200-463) did not interact with the Oskar-eLOTUS domain, and similarly, neither the unstructured region of Oskar (Oskar 241-387) nor its OSK domain (Oskar 388-606) interacted with Vasa-CTD (**Fig. 1c)**. Surface point mutations that had been previously shown to interfere with the Vasa-Oskar interaction ^28^ were also found to be inhibitory in the ReLo assay (**Fig. 1d**). Together, these data confirmed the specific interaction between Vasa-CTD and Oskar-eLOTUS and demonstrated that ReLo can be applied to map PPIs and to identify mutations that interfere with PPIs.

Oskar-eLOTUS and Vasa-CTD form a transient complex characterized by a dissociation constant (K_D_) of ∼10 µM, and although this complex has been crystallized, it is not sufficiently stable to be detected by size-exclusion chromatography ^28, 30, 31^. Nevertheless, the relocalization upon the interaction between Oskar-eLOTUS and Vasa-CTD was clearly detectable in the ReLo assay: in a total of three independent replicates, the relocalization of Vasa-CTD toward the plasma membrane was observed in 91 of 94 (i.e., 97%) cotransfected cells with both red and green fluorescent signals (**Supplementary Fig. 2**). These data indicate that relocalization is a highly frequent event and that the assay is well suited for studying low-affinity complexes. Most other interactions tested revealed relocalization in 100% of the cotransfected cells (see below; **Supplementary Fig. 3**).

### Conformation-dependent interactions

Previous data have suggested that the Oskar-Vasa interaction depends on the conformation of Vasa ^28^. We aimed to test the interaction between Oskar and different Vasa conformations in a full-length protein context. However, in contrast to the eLOTUS domain of Oskar, full-length Short Oskar did not localize to the cytoplasm but exclusively localized to the nucleus in S2R+ cells (**Supplementary Fig. 1b)** ^28^. Therefore, we made use of the OST4-mCherry-Oskar construct, which localized to membranous structures within the cytoplasm (**Supplementary Fig. 1d and Fig. 1e**). Upon coexpression, wild-type Vasa relocalized and colocalized with OST4-Oskar at the ER (**Fig. 1e**), confirming the interaction between Vasa and Oskar in the ReLo assay. Also using the OST4 anchor, the interaction mapped to the Oskar-eLOTUS domain and the Vasa-CTD **(Supplementary Fig. 1e**) and the known Vasa interface mutant (F504E) was unable to bind to Oskar (**Fig. 1e**), which is consistent with the data obtained using the PH domain (**Fig. 1c, 1d**).

The cores of Vasa and other ATP-dependent DEAD-box RNA helicases are composed of two RecA-like domains, which display different orientations relative to each other depending on the presence of bound ATP and RNA ^32^. In a substrate-unbound form, the helicase core adopts an open conformation and closes upon substrate binding (**Fig. 2a)**. To assess the conformation-dependent Vasa interaction with the ReLo assay, we used Vasa variants with well-characterized point mutations in the ATP-binding pocket that stabilize either the open conformation (K295N; Vasa-open) or the closed conformation (E400Q; Vasa-closed) ^33–35^. When testing the interaction of OST4-anchored Oskar with the Vasa mutants using the ReLo assay, we observed an interaction with Vasa-open but not Vasa-closed (**Fig. 2b)**, which revealed that Oskar prefers to bind to the open conformation of Vasa. These outcomes were consistent with our previous observations ^28^ and demonstrated that the ReLo assay allows the study of PPIs that are dependent on the specific conformation of an interaction partner.

**Fig. 2.**
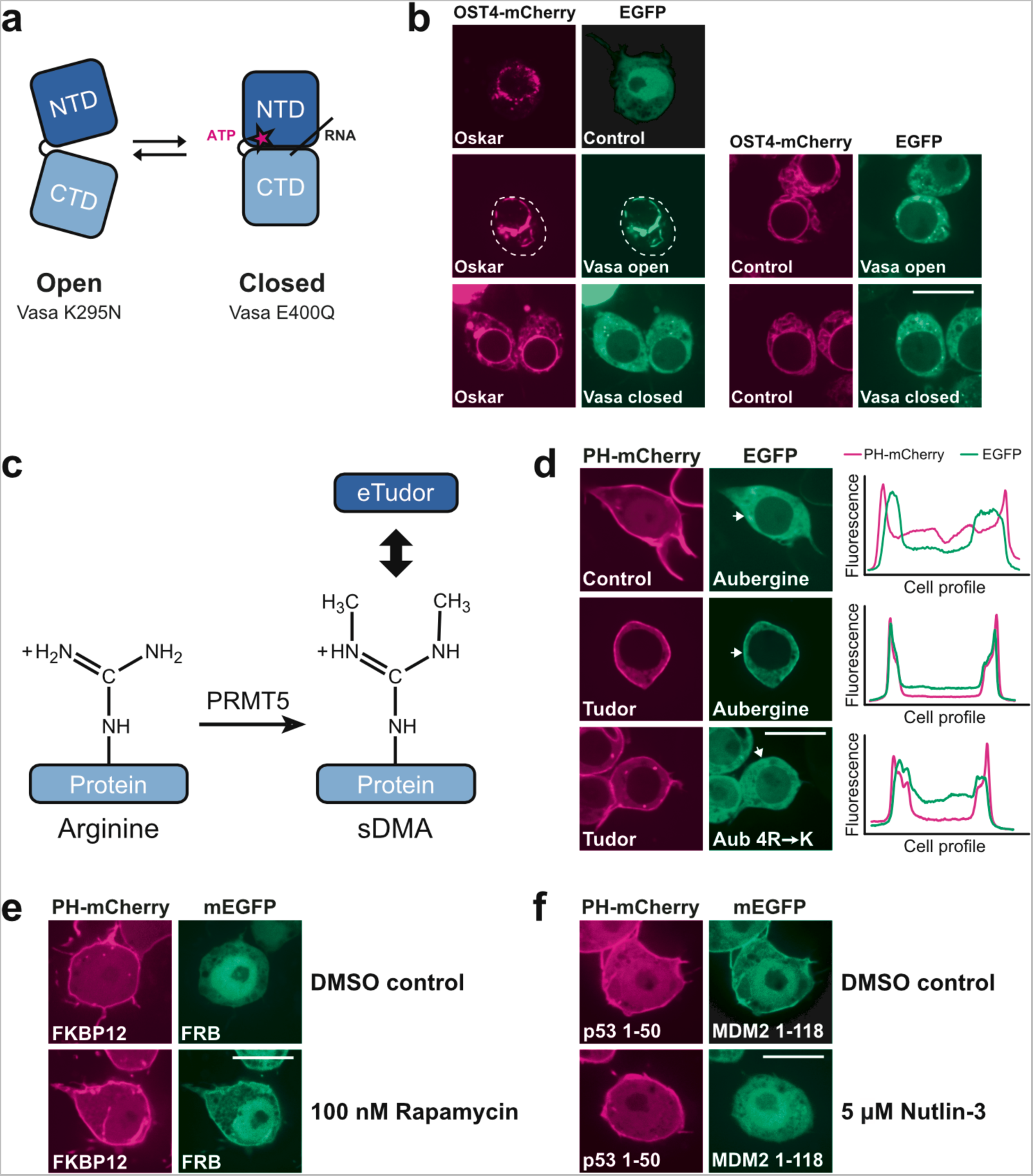
PPI studies using ReLo with respect to conformation, posttranslational modification and drug sensitivity. **a** Pictorial representation of Vasa N- and C-terminal RecA-like domains (NTD and CTD, respectively) in the open and closed conformations. Vasa-open and Vasa-closed carry the K295N or E400Q mutations, respectively. **b** EGFP-Vasa-open, but not EGFP-Vasa-closed, interacted with OST4-mCherry-Oskar. The white dotted lines indicate cell boundaries. **c** Symmetric dimethyl arginine (sDMA) modifications can be catalyzed by PRMT5 and recognized by eTud domains. **d** Wild-type Aub interacted with Tudor, whereas Aub carrying nonmethylatable 4R->K point mutations (R11K/R13K/R15K/R17K) did not. The profiles indicate the measured fluorescence distribution across the cell along a line indicated by the white arrows (shown for the green channel only). Red and green fluorescence signals are independent and displayed relatively. **e** Rapamycin induced an interaction between human FKBP12 and FRB. The control contained 0.0009% dimethyl sulfoxide (DMSO). **f** The interaction between p53 1-50 and MDM2 1-118 was inhibited by nutlin-3 treatment. The control contained 0.05% DMSO. In the “Control” experiments the respective construct was coexpressed with the vector indicated at the top of the panel lacking an insert. The scale bar is 10 µm.

### Protein arginine methylation-dependent interactions

Protein interactions may depend on posttranslational modifications, such as protein arginine methylation. The symmetric dimethylated arginine (sDMA) modification is catalyzed by a subset of protein arginine methyltransferases (PRMTs) ^36^, and sDMA methylation activity has been previously reported in S2 cells ^37^. To test whether the ReLo assay is suitable for investigating interactions involving sDMA modifications in S2R+ cells, we tested the previously characterized strictly sDMA-dependent interaction between the PIWI protein Aubergine (Aub) and Tudor ^38, 39^ (**Fig. 2c)**. Using ReLo, we indeed observed an Aub-Tudor interaction (**Fig. 2d**). Aub carries four sDMAs within its RG-rich N-terminus (R11, R13, R15, and R17), which are specifically recognized and bound by extended Tudor (eTud) domains of Tudor ^38–41^. The substitution of these four arginine residues with lysine residues (Aub R->K) rendered Aub unmodifiable by PRMT5 ^41^ and abolished the Aub-Tudor interaction (**Fig. 2d**). Together, these data demonstrated that the S2R+ cells exhibited sufficient sDMA activity to effectively modify proteins expressed after transient transfection of the cells. We conclude that ReLo is suitable for the study of PPIs that depend on PRMT5-catalyzed sDMA modification.

### Effect of small molecules on PPIs

We subsequently tested whether the ReLo assay can be used to study PPIs that are induced by the addition of small molecules to the cell culture medium. To this end, we tested the previously characterized rapamycin-dependent interaction between human FK506-binding protein 12 (FKBP12) and the FKBP12-rapamycin-binding domain (FRB) of human mTOR ^42^. In the absence of rapamycin, using the dimethyl sulfoxide (DMSO)-containing control medium, FKBP12 and FRB did not interact in the ReLo assay but did interact in the presence of 100 nM rapamycin in the cell culture medium (**Fig. 2e**).

Furthermore, we tested whether the ReLo assay can be used to inhibit PPIs through drug treatment. We opted to interfere with the interaction between human p53 and the human ortholog of mouse double minute 2 (MDM2) using the known peptidomimetic inhibitor nutlin-3 ^43^. In the ReLo assay, we used only the N-terminal domains of p53 and MDM2, which were sufficient to mediate the p53-MDM2 interaction ^44^. Indeed, although an interaction between p53 1-50 and MDM2 1-118 was detected in the DMSO control experiment, this interaction was not observed when the cells were incubated with 5 µM nutlin-3 (**Fig. 2f**).

Together, these data suggest that the ReLo assay can be used to study interactions involving non-*Drosophila* proteins and to screen drugs based on their ability to either enable or inhibit a specific PPI.

### ReLo reveals direct binary interactions

Thus far, we have tested known direct interactions between proteins that are not endogenously expressed in S2R+ cells. PPIs detected with the ReLo assay may not necessarily involve direct contact. Instead, PPIs may also result from indirect contacts caused by incorporation of the coexpressed pair of proteins into cell-endogenous protein complexes and subsequent indirect bridging of this pair through one or more common interaction partners. To understand the extent to which direct or indirect associations underlie a relocalization event observed with ReLo, we assessed interactions between individual subunits of the CCR4-NOT complex, which is endogenous to S2R+ cells. The CCR4-NOT complex is an essential eukaryotic deadenylase comprising six subunits that form the core with an architecture that is well-characterized at the molecular and structural levels ^45, 46^. In this complex, NOT1, which has a size of 281 kDa in *Drosophila,* acts as the scaffolding subunit for the assembly of all other subunits. Specifically, the CAF1-CCR4 heterodimer and CAF40 bind to the central region of NOT1, and the NOT2-NOT3 subcomplex associates with the C-terminal region of NOT1 (**Fig. 3a)**. Using ReLo, we performed a systematic pairwise screen in which each subunit was tested against all other subunits of the *Drosophila* CCR4-NOT core complex (**Fig. 3b and Supplementary Fig. 4a)**. In S2R+ cells, NOT1 and NOT3 localized exclusively to the cytoplasm, whereas NOT2, CAF1, CAF40 and CCR4 localized to both the cyto- and the nucleoplasm (**Supplementary Fig. 4b)**. Remarkably, we observed interactions only between proteins that had previously been shown to exhibit direct associations but not between indirectly linked combinations: NOT1 bound specifically to CAF1, CAF40, NOT2, and NOT3, while CAF1 bound to CCR4, and NOT2 bound to NOT3 (**Fig. 3b, c, Supplementary Fig. 4a, b)**. We also tested the interactions to CAF1 using the OST4 membrane anchor and observed results similar to those obtained using the PH anchor (**Supplementary Fig. 4c)**. Together, these data suggest that the incorporation of two tagged CCR4-NOT complex subunits into one endogenous complex is insignificant, and thus direct but not indirect interactions between the two proteins tested are observed with the ReLo assay.

**Fig. 3.**
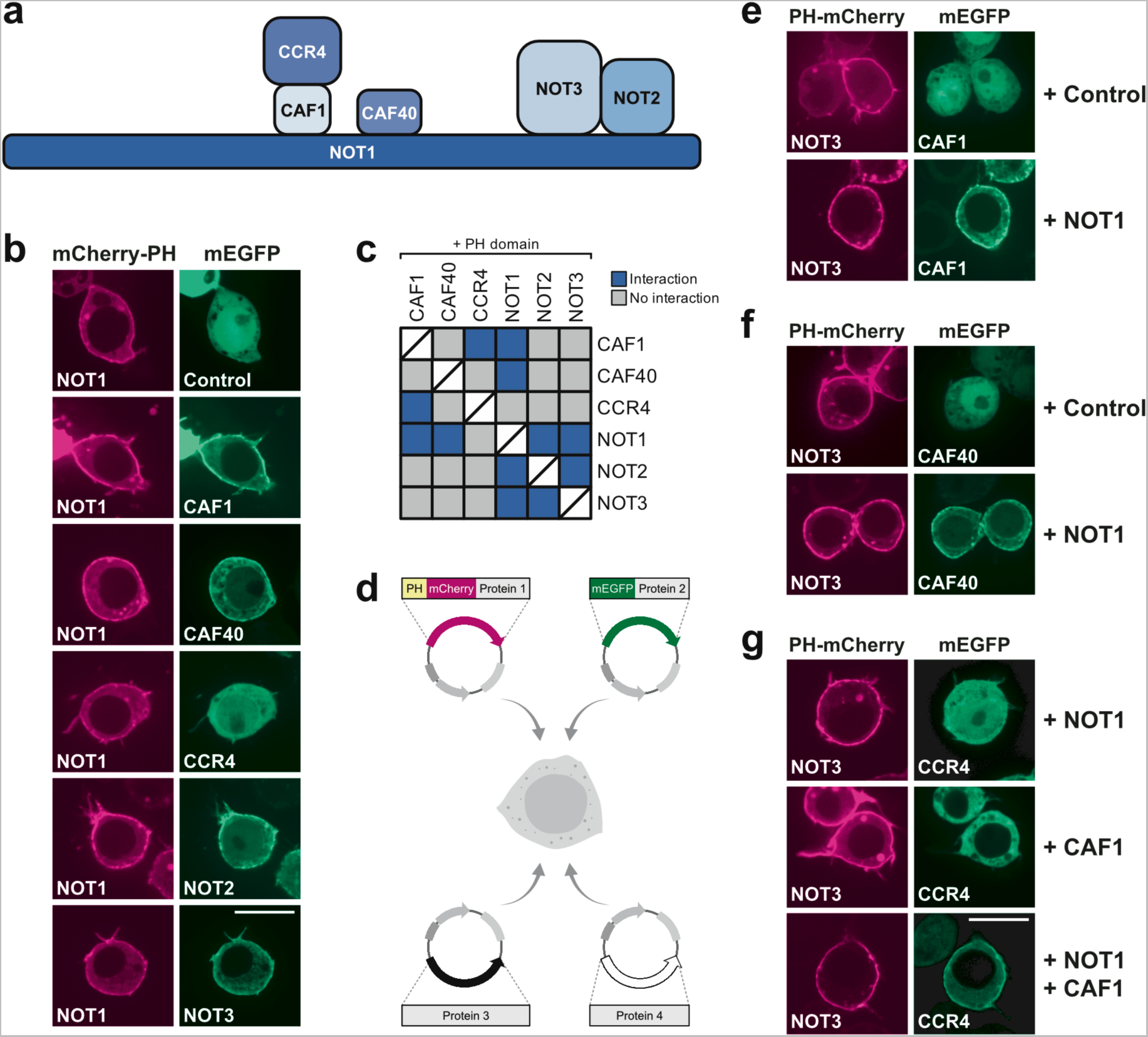
ReLo identifies direct PPIs. **a** Subunit organization of the *Drosophila* CCR4-NOT core complex. **b** NOT1-mCherry-PH recruited CAF1, CAF40, NOT2, and NOT3 but not CCR4 to the plasma membrane. See **Supplementary Fig. 3a** for the PPI analysis of the other CCR4-NOT complex subunits. **c** Summary of the results from a pairwise screen of CCR4-NOT complex core subunits using ReLo. The blue color indicates an interaction, and the gray color indicates no interaction. **d** Schematic showing the ReLo assay involving coexpression of four different protein constructs. **e** NOT3 interacted with CAF1 in the presence of NOT1. **f** NOT3 interacted with CAF40 in the presence of NOT1. **g** NOT3 recruited CCR4 to the plasma membrane only in the presence of both NOT1 and CAF1. In the “Control” experiments the respective construct was coexpressed with the vector indicated at the top of the panel lacking an insert. The scale bar is 10 µm.

Although the subunits of the endogenous CCR4-NOT complex show moderate to high expression levels in S2R+ cells ^47^ (**Supplementary Fig. 5a)**, we aimed to challenge the ReLo assay by testing PPI between subunits of a cytoplasmic complex that is even more abundant in S2R+ cells. We chose the seven-subunit Arp2/3 complex, which is known to initiate actin polymerization in eukaryotes ^48^, and of which several subunits are among the most highly expressed cytoskeletal genes ^47, 49^ (**Supplementary Fig. 5a)**. In S2R+ cells, the Arp2/3 subunits Arp3, Arpc1, and Arpc4 are expressed about two- to four-fold higher as compared to NOT3, which is the most abundant subunit of the CCR4-NOT complex (**Supplementary Fig. 5a)**. Compared to the CCR4-NOT complex, binary PPIs within the Arp2/3 complex are less well characterized, and we concluded which subunit is directly or indirectly bound to another subunit based on the crystal structure of the bovine Arp2/3 complex ^50^ (**Supplementary Fig. 5b, c)**. The crystal structure points to six direct pairwise contacts. Using ReLo assays, we detected two PPIs, namely the Arpc2-Arpc4 and the Arpc4-Arpc5 interaction (**Supplementary Fig. 5c, d**). The other four contacts, which we did not observe with ReLo, were also not detected in an earlier pairwise study of the human Arp2/3 complex using Y2H assays ^51^. Importantly, we did not detect any false-positive interaction using the ReLo assay. These data suggest that even if a complex is highly abundant in a cell, the detection of a PPI between two protein candidates in the ReLo assays is direct rather than indirectly mediated by incorporation into the same endogenous complex. Nevertheless, we cannot exclude that occasionally, indirect interactions are detected through a naturally expressed bridging protein.

Finally, we compared the ReLo assay to the split-ubiquitin yeast two-hybrid assay ^27, 52^, which we have previously used successfully to map the interaction between Short Oskar and Vasa ^30^. When we assessed the pairwise interactions between the six CCR4-NOT complex core subunits using Y2H, we observed mostly false negative and false positive results; the only conclusive interaction that we detected was between NOT2 and NOT3 (**Supplementary Fig. 6**). This observation is in stark contrast to our data obtained using ReLo assays. As both Y2H and ReLo worked well to map the Oskar-Vasa interaction, together the data suggest that the applicability of Y2H for testing interactions is case/project dependent. ReLo could be expected to be more reliable than Y2H for testing large proteins from higher eukaryotes, although a larger scale study would be required to fully test this.

### Topological description of multisubunit complexes

As has been described for S2 cells ^53^, we usually observed a very high cotransfection efficiency of the S2R+ cells when performing ReLo experiments. Therefore, we tested whether the bridging of CCR4-NOT subunits that do not directly interact is observable when one or two common binding partners are added to the mixture. Specifically, S2R+ cells were cotransfected with three or four plasmids: one plasmid expressing the PH-mCherry fusion construct, one plasmid expressing the mEGFP fusion construct, and one or two plasmids expressing nonfluorescent bridging factors (**Fig. 3d**). Indeed, NOT3 failed to interact with CAF1 or CAF40 in the presence of the control plasmid but interacted when NOT1 was provided in addition (**Fig. 3e, f**). We also tested the bridging of the NOT3-CCR4 interaction, which requires not only NOT1 but also the CAF1 subunit. The NOT3-CCR4 interaction was not observed if NOT1 alone was coexpressed but was observed when both NOT1 and CAF1 were coexpressed (**Fig. 3g**). Similar results were obtained from the testing of indirect interactions with CAF40 (**Supplementary Fig. 4d**). Together, these data show that bridging experiments with ReLo can be used to reconstitute the binding topology of multisubunit complexes such as the CCR4-NOT complex.

### Assessment of PPIs between the CCR4-NOT complex and mRNA repressor proteins

The CCR4-NOT complex can be recruited specifically to mRNAs through adapter RNA-binding proteins, which leads to accelerated deadenylation and eventually degradation and/or translational repression of the targeted mRNA. We then assessed PPIs between some of these repressor proteins and the six core subunits of the CCR4-NOT complex using ReLo (**Fig. 4a)**. Co-IP experiments combined with structural analysis previously revealed the specific subunit(s) of the CCR4-NOT complex through which adapter proteins recruit the complex to a specific mRNA. For example, *Drosophila* Bag-of-marbles (Bam) has been shown to bind specifically to CAF40 ^54^. We confirmed this finding with ReLo: Bam bound to CAF40 but not to any other CCR4-NOT complex subunit (**Fig. 4b and Supplementary Fig. 7a)**. A point mutation in Bam (M24E; Bam MUT), which is known to interfere with CAF40 binding ^54^, also abolished CAF40 binding in the ReLo assay (**Fig. 4b)**. Likewise, a mutation at the CAF40 surface (V186E; CAF40 MUT), which was shown to prevent Bam binding ^54^, abolished the interaction with Bam in the ReLo assay (**Fig. 4b)**.^55^

**Fig. 4.**
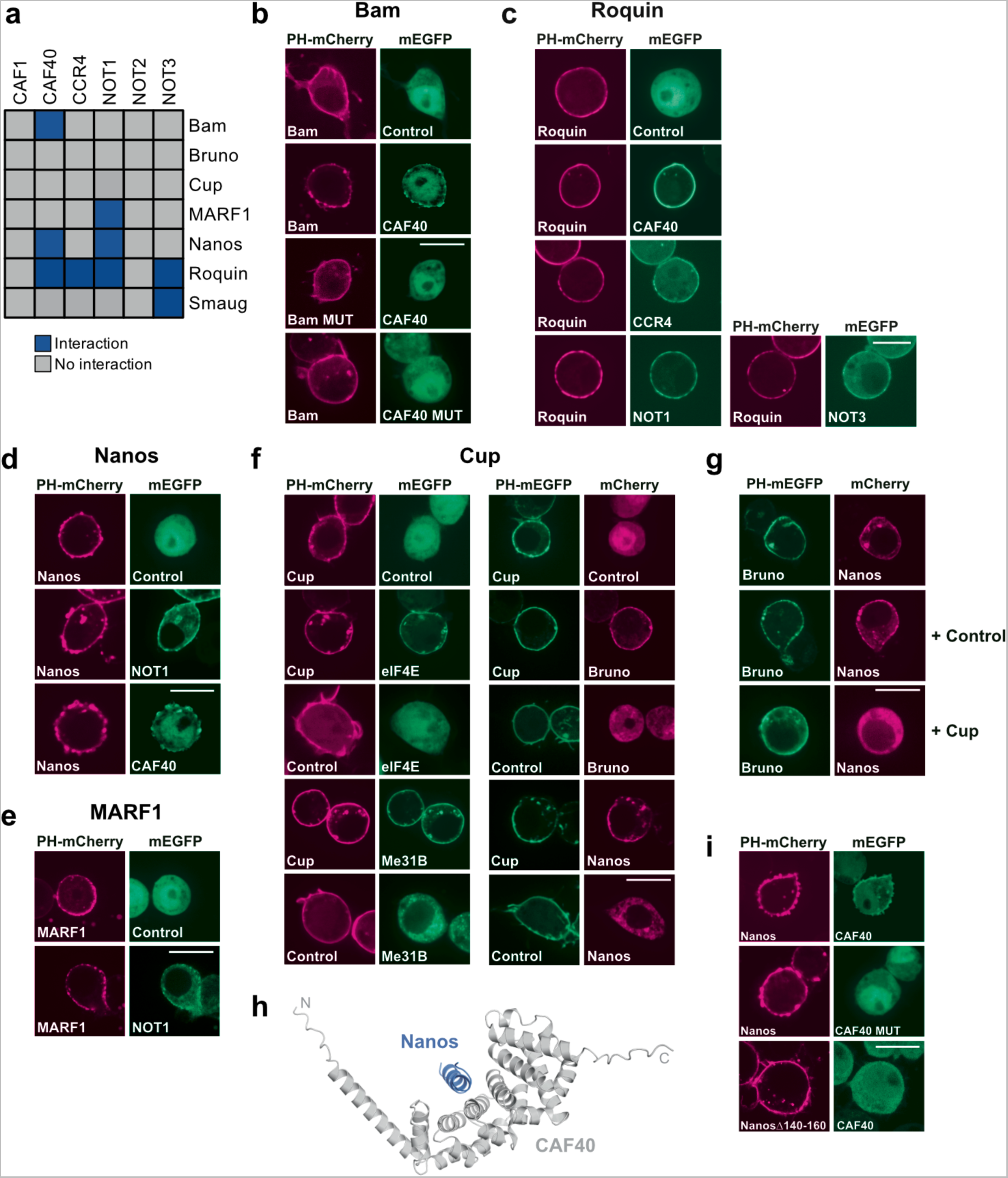
PPIs between the CCR4-NOT complex and repressor proteins. Summary of the results (**a**) from a pairwise screen of CCR4-NOT complex core subunits with Bam (**b**), Nanos (**c**), Roquin (**d**), MARF1 (**e**), Bruno and Cup (**Supplementary Fig. 7**), and Smaug ^63^. The blue color indicates an interaction, and the gray color indicates no interaction. Bam MUT carries the M24E and CAF40 MUT the V186E point mutation. **f** Cup interacted with eIF4E, Me31B, Bruno, and Nanos. **g** Bruno interacted with Nanos in the absence but not in the presence of Cup. In the “Control” experiments the respective construct was coexpressed with the vector indicated at the top of the panel lacking an insert. The scale bar is 10 µm.

*Drosophila* Roquin has been demonstrated to bind to *Drosophila* CAF40. It also binds to the so-called CAF40 module composed of CAF40 and the CAF40-binding region of NOT1 and to the NOT1/2/3 module composed of the C-terminal domains of the NOT1, NOT2, and NOT3 subunits of human CCR4-NOT complex ^55^. Of the subunits tested in the ReLo assay, we detected clear binding of Roquin to *Drosophila* CAF40, NOT1, and NOT3 but not to NOT2 (**Fig. 4c and Supplementary Fig. 7b)**, essentially confirming the previous data. In addition, we detected a previously unknown interaction between Roquin and CCR4 (**Fig. 4c**).

*Drosophila* Nanos was shown to bind to the NOT1/2/3 module of the human CCR4-NOT complex in GST pull-down experiments, but does not bind to any of these domains individually ^56^. With ReLo, we detected the interaction of Nanos with *Drosophila* NOT1 but not with NOT2 or NOT3, which suggests that the NOT1 interaction might be predominant (**Fig. 4d and Supplementary Fig. 7c)**. In addition, we detected Nanos binding to the CAF40 subunit (**Fig. 4d)**, an interaction that has not previously been reported.

Encouraged by these mostly validating data, we tested the interaction of the CCR4-NOT complex with less well characterized potential adaptor proteins. *Drosophila* meiosis regulator and mRNA stability factor 1 (MARF1) is an oocyte-specific protein that recruits the CCR4-NOT complex to target mRNAs and thereby controls meiosis ^57^. *Drosophila* Smaug recruits the CCR4-NOT complex to *nanos* and other mRNAs to regulate posterior patterning and nuclear divisions of the early embryo^58–62^. Whether MARF1 or Smaug directly associate with the CCR4-NOT complex was unknown. Using ReLo, we found that Smaug interacted with the NOT3 subunit ^63^, and that MARF1 bound to the NOT1 subunit (**Fig. 4e and Supplementary Fig. 7d**), which suggests that Smaug and MARF1 are indeed direct recruiters of the CCR4-NOT complex.

*Drosophila* Cup and its orthologous protein 4E-T are translation repressor proteins that act through their interactions with the eukaryotic translation initiation factor 4E (eIF4E) and the DEAD-box RNA helicase Me31B/DDX6 ^64–68^. Both the human 4E-T - Me31B and the *Drosophila* Cup-eIF4E complex are structurally characterized ^69, 70^ and we detected both interactions using ReLo assays (**Fig. 4f, left panels**). As demonstrated through co-IP experiments, Cup also interacts with the CAF1, CCR4, NOT1, NOT2, and NOT3 subunits of the CCR4-NOT complex ^71^. We did not detect Cup binding to any subunit of the CCR4-NOT core complex using ReLo assays (**Supplementary Fig. 7e**), which suggests that if Cup directly recruits the CCR4-NOT complex, then it probably involves weak and multivalent interactions. Cup also reportedly binds the repressors Nanos and Bruno ^65, 72, 73^, and with the ReLo assay we confirmed both interactions (**Fig. 4f, right panels**). Consistent with a recent report ^74^, we also observed that Bruno binds directly to Nanos (**Fig. 4g**). We then asked if Bruno and Nanos bind to Cup as a complex. However, when coexpressed with Cup, Bruno and Nanos did not interact (**Fig. 4g**), which suggests that Bruno and Nanos compete for binding to the same region of Cup. Bruno has thus far not been shown to establish direct contact with the CCR4-NOT complex, and we did not detect Bruno binding to any of the core CCR4-NOT complex subunits (**Supplementary Fig. 7f**).

In summary, the assessment of PPIs using ReLo involving a wide selection of repressor proteins, confirmed many previously described interactions and revealed new interactions. Thus, ReLo is a powerful method for identifying novel PPIs in a candidate approach.

### Combining ReLo assays with structural prediction

Lastly, we aimed to obtain some molecular insight into one of the newly identified interactions. We tested the structural prediction of the Nanos-CAF40 complex using AlphaFold2-multimer-v3 and obtained a high-confidence model as indicated by the predicted aligned error (PAE) plot (**Fig. 4h, Supplementary Fig. 8a**) ^75–77^. The predicted structure is composed of full-length CAF40 and a short α-helix spanning residues 140 to 160 of Nanos, which is bound to the same site of CAF40 that was shown to accommodate the Bam or Roquin α-helical peptide ^54, 55^ (**Supplementary Fig. 8b**). The CAF40 point mutation that prevented Bam binding (**Fig. 4b**), also abolished the interaction with Nanos (**Fig. 4i**). Likewise, deletion of the CAF40-binding α-helix from Nanos (Nanos τι140-160) prevented the interaction with CAF40 (**Fig. 4i**). These data are consistent with the predicted structural model of the Nanos-CAF40 complex. Taken together, our exemplary analysis suggests that the combination of ReLo assays with structural modeling can lead to fast molecular insight into protein complexes.

## DISCUSSION

The detection and characterization of PPIs are crucial for uncovering regulatory mechanisms that underlie cellular processes. Here, we described ReLo, a rapid procedure for identifying and investigating pairwise as well as multisubunit PPIs. We provided strong evidence showing that pairwise PPIs identified with ReLo are based on direct contacts. Thus, PPI partners newly identified using ReLo are highly promising candidates for use in subsequent studies using *in vitro* assays and experimental structural biology methods. Alternatively, PPI mapping data obtained from ReLo experiments may be used to guide subsequent modeling experiments using AlphaFold2-Multimer to obtain structural information on complexes ^75, 76^. A thus predicted protein–protein interface may then rapidly be validated through mutational analysis using ReLo, and the mutations that specifically interfere with a PPI may finally be tested *in vivo* without the need for purifying a protein or experimentally determining a structure of a protein complex.

We refrained from quantifying the relocalization events observed with ReLo through image processing tools to assess or compare PPIs. Although only qualitative, the results obtained were typically clear. For example, in control experiments, we never observed relocalization, and in experiments in which proteins interacted, most or all cells showed relocalization (**Supplementary Figs. 2 and 3**). Furthermore, the implementation of a relocalization score as a quantified readout of an interaction may have easily led to the false assumption that the score represents a measure of binding affinity between proteins. Such false assumptions are also an issue with Y2H dot assays, which appear to but do not provide quantitative data. The degree of relocalization identified with the ReLo assay or yeast cell growth in a Y2H assay not only reflects the binding affinity of the protein complex but also relies on the protein expression levels, protein stability, subcellular localization, and other factors. With this thought in mind, we prefer to consider ReLo a qualitative PPI method, the strength of which lies in its speed and versatility.

ReLo is easy to perform in every laboratory equipped with cell culture technology and with access to a confocal fluorescence microscope. In contrast to standard Y2H or PCA assays, the expression levels of the proteins tested in a ReLo assay are directly monitored during microscopy, which facilitates data interpretation. The *Drosophila* S2R+ cell line used for our ReLo tests requires very simple handling. Similar to S2 cells, S2R+ cells grow at ambient temperature, do not need an incubator with CO_2_, and can be passaged without the need for coated dishes nor the scraping or trypsinization of cells ^53^. Using S2R+ cells, we successfully investigated not only *Drosophila* PPIs but also human PPIs. However, when a cell line derived from an alternative organism is needed for a ReLo assay, only the PH or OST4 membrane anchors need to be inserted into expression vectors compatible with the desired cell line.

With ReLo assays, we investigated PPIs involving NOT1 and Tudor, two large proteins comprising 281 kDa (2505 aa) and 285 kDa (2515 aa), respectively. Thus, the detection of PPIs using the ReLo assay appears to be successful regardless of the length of the protein of interest, which, in our experience, is a big advantage over Y2H methods. Similar to any other cell-based PPI assay, attention is needed when working with toxic proteins. To reduce toxicity, we recommend testing the splitting of the toxic protein into its domains or, when possible, testing protein variants with mutated active sites.

PPIs are highly relevant as putative therapeutic targets for the development of new treatments ^78, 79^. In ReLo, complex formation is reversible, and we demonstrated that ReLo is a tool for testing the effect of small molecules on PPIs. Due to its simple setup, ReLo might be easily adjusted to a high-throughput approach using automated imaging, which allows for large drug screening experiments. In a setup in which a single specific PPI is subjected to a drug screening experiment, it might be advantageous to express the two protein partners from one plasmid to ensure equimolar protein expression within a cell, which may facilitate data interpretation.

Taken together, our data show that investigations with ReLo are rapid, simple, and reliable. We recommend using ReLo as an initial tool to screen and characterize PPIs, particularly in cases in which Y2H or more complicated approaches fail. Subsequently, ReLo can be complemented with biochemical, structural, or genetic approaches to further characterize or ultimately validate the biological relevance of a given PPI.

## ACKNOWLEDGMENTS

We thank Elmar Wahle, Julien Bethune, Bernd Bukau, and Ivana Vonkova for DNA constructs and Aurelio Telemann for the S2R+ cell line. We are grateful to Christian Bleischwitz, Eva Boberlin, Jana Kubíková, Katharina Müller, Rebecca Reinig, Gabrielė Ubartaitė, and Xiaohan Zhao for technical assistance provided with some experiments. We thank the Nikon Imaging Center at the University of Heidelberg for access to microscopes. We gratefully acknowledge the data storage service SDS@hd supported by the Ministry of Science, Research and the Arts Baden-Württemberg (MWK) and the German Research Foundation (DFG) through the grants INST 35/1314-1 FUGG and INST 35/1503-1 FUGG. We thank Peter Becker, Hüseyin Besir, Julien Béthune, and Doris Höglinger for their insightful comments on the manuscript. This work was funded by the Emmy-Noether Program of the German Research Foundation (DFG; JE-827/1- 1).

## METHODS

### Plasmid backbone construction

pAc5.1-EGFP and pAc5.1-mCherry were described previously ^28^. pAc5.1-mEGFP encodes the monomeric A206K EGFP variant and was generated by site-directed mutagenesis of pAc5.1-EGFP. The pAc5.1-λN-HA vector ^80^ was used to create nonfluorescent constructs. The *Saccharomyces cerevisiae* OST4 sequence was amplified from a pDHB1 vector ^27^ and inserted into the KpnI site of pAc5.1-mCherry to yield pAc5.1-OST4-mCherry. For cloning into the pAc5.1-EGFP, pAc5.1-mEGFP, pAc5.1-mCherry, pAc5.1-λN-HA, and pAc5.1-OST4-mCherry vectors, sequences of interest were inserted into the blunt-end EcoRV site. The rat PLCδ_1_-PH sequence was amplified from the pETM11-His6-PH-Sumo3-sfGFP vector ^81^ and inserted into the KpnI site of pAc5.1-mCherry and pAc5.1-mEGFP to obtain pAc5.1-PH-mCherry and pAc5.1-PH-mEGFP, respectively. Alternatively, the PH sequence was inserted into the EcoRV site of pAc5.1-mCherry to obtain the pAc5.1-mCherry-PH vector. For all PH-containing vectors, a new unique in-frame FspAI blunt end site was introduced 3’ or 5’ with respect to the fluorescent protein sequences and was used to insert the sequence of interest.

### Cloning

Ligation reactions were assembled in a 10 µl reaction containing T4 DNA ligase (Thermo Scientific), 50 ng of vector DNA, and a 5- to 20-fold molar excess of DNA inserts and were incubated for 1 to 2 h at room temperature. DNA inserts were generated by PCR amplification. To prevent re-ligation, the reaction was supplemented with 0.25 µl of the blunt end restriction enzyme that was also used for linearization of the respective vector, except when this site was present in the insert sequence. Positive clones were screened by colony PCR using one primer that binds to the vector and a second that binds to the insert. All constructs were verified by sequencing. Detailed information on all DNA constructs used in this study is listed in **Supplementary Table 1**.

### Cell culture

S2R+ cells were cultured at 25°C in Schneider’s *Drosophila* medium + (L)-glutamine (Thermo Scientific) supplemented with 10% fetal bovine serum (Sigma) and 1 x Gibco™ Antibiotic-Antimycotic (Thermo Scientific). Cells were seeded in six-well glass-bottom plates (Cellvis; earlier experiments) or four-well polymer µ-slides (Ibidi; later experiments) and cotransfected using jetOPTIMUS (Polyplus Transfection) according to the instruction manual. Specifically, 600 µl of cells with a density of 1×10^6^ cells/ml were seeded per well in a four-well slide and 61 µl of transfection reagent mixture was added. This mixture contained 1 µl of transfection reagent and 600 ng of DNA in total for cotransfecting two (300 ng DNA each), three (200 ng each), or four (150 ng each) plasmids diluted in jetOPTIMUS buffer. After 48 h of incubation at 25°C, images of live cells were taken with a 60x or 100x oil objective and a Nikon TE2000 laser scanning, a Nikon Ti-E spinning disc confocal, or a Nikon Eclipse Ti2 laser scanning fluorescence microscope and processed with Fiji software ^82^. Changing the amount of DNA or transfection reagent during the transfection did not change the protein expression levels in the cells. However, reducing the total DNA amount to 150 ng and/or the transfection reagent to 0.5 µl significantly reduced the transfection efficiency.

### Drug treatment

Rapamycin and nutlin-3 (both from MedChemExpress) were dissolved in DMSO to obtain stock concentrations of 10.9 mM and 10 mM, respectively, and they were diluted with SF-4 Baculo express medium (BioConcept) to generate working concentrations of 100 nM and 5 μM, respectively. S2R+ cells were grown in SF-4 medium and cotransfected with desired plasmids using FuGENE® HD transfection reagent (Promega). Specifically, cells were seeded as described above and 26 µl of transfection reagent mixture was added. This mixture contained 1 µl of transfection reagent and 400 ng of DNA (i.e., 200 ng DNA each for cotransfecting two plasmids) diluted in SF-4 Baculo express medium. Twenty-four hours after transfection, the cells were treated with the drug by replacing the medium with drug-containing medium. Cells were imaged after 24 h of incubation with the drug or DMSO control medium at 25°C.

## AUTHOR CONTRIBUTIONS

H.K.S. and J.M. acquired, analyzed, and interpreted the data. M.J. conceived and supervised the project, acquired, analyzed, and interpreted the data, and wrote the manuscript with input from all authors.

## DATA AVAILABILITY STATEMENT

The authors declare that all relevant data supporting the findings of this study are available within the article and its supplementary information files.

