## Supplementary Data for "ReLo is a simple and quick colocalization assay to identify and characterize direct protein-protein interactions"

Inventory:

Supplementary Figures 1-8

Supplementary Tables 1

Supplementary Methods

Supplementary References

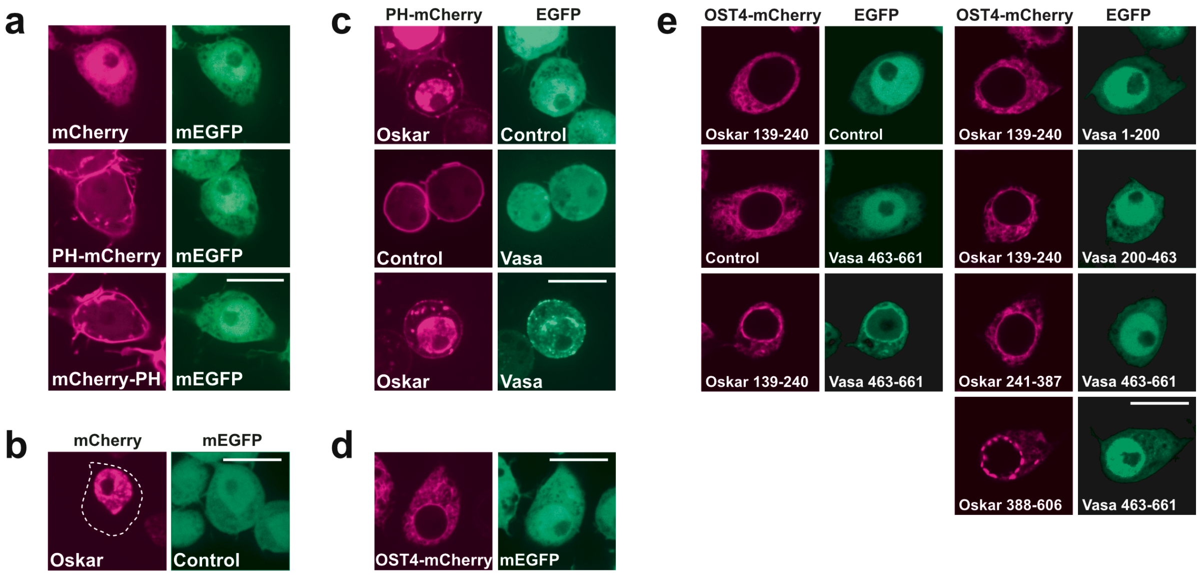


**Supplementary Fig. 1. ReLo using the PH domain or the OST4 fusion.**
mCherry, PH-mCherry, OST4-mCherry, or mEGFP alone or as fusions to the proteins indicated were coexpressed in S2R+ cells and their localization was analyzed by microscopy. **a** Both N- and C-terminal fusions to the PH domain directed the localization of mCherry to the plasma membrane. **b** mCherry-Oskar localized to the nucleus. **c** PH-mCherry Oskar retained in the nucleus and only partially localized to the plasma membrane (top panel). Vasa only partially relocalized with Oskar to the plasma membrane (bottom panel). **d** A fusion to OST4 directed the localization of mCherry to the ER. **e** Vasa 463-661 but not Vasa 1-200 or 200-463 interacted with the Oskar eLOTUS domain (139-249). Vasa 463-661 did not interact with Oskar 241-396 or Oskar 398-606. The scale bar is 10 µm.

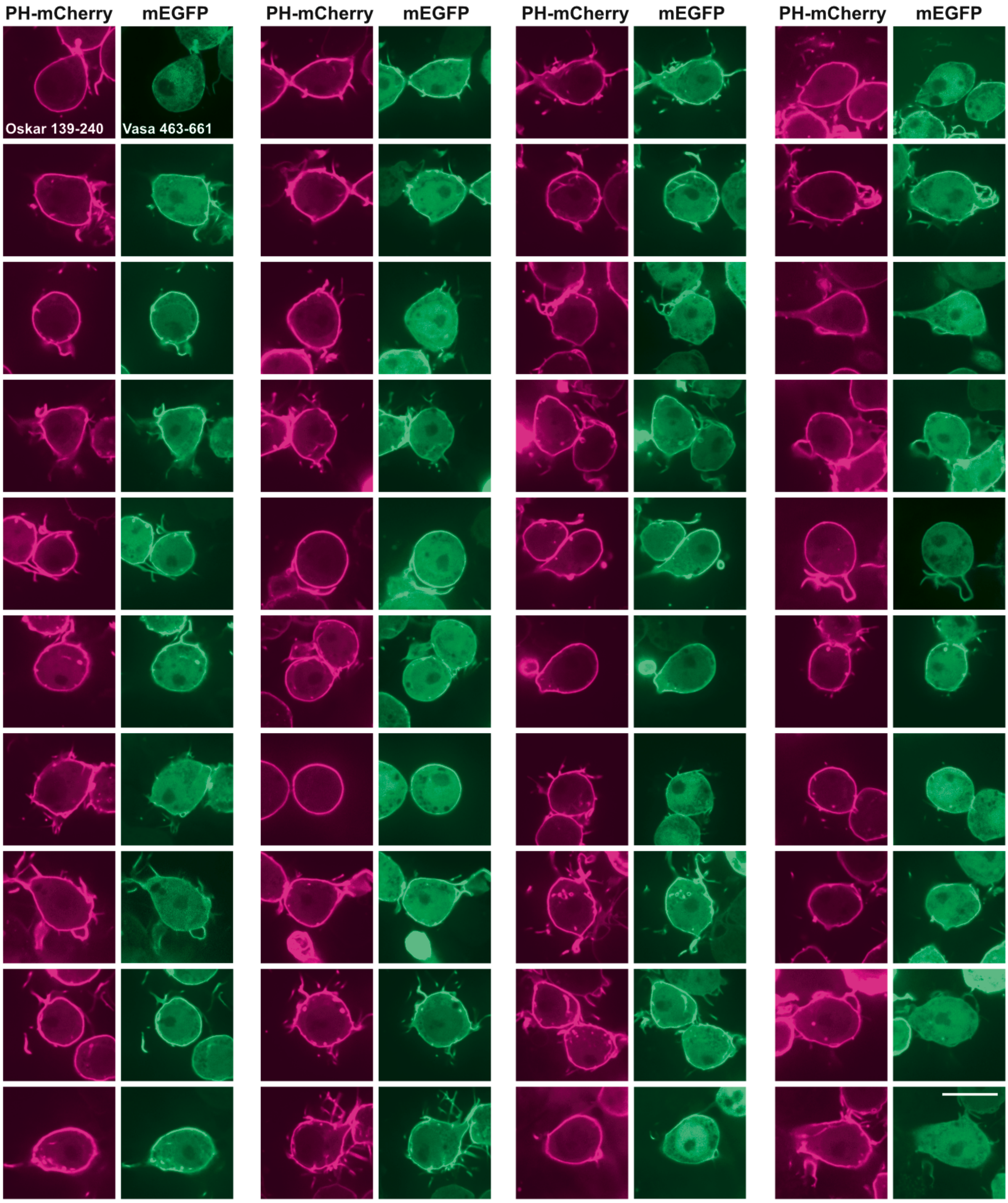


**Supplementary Fig. 2. Assessing the Oskar-eLOTUS - Vasa-CTD interaction.** PH-mCherry-Oskar 139-240 and mEGFP-Vasa 463-661 were coexpressed in S2R+ cells and their localization was analyzed by microscopy. In 38 of 40 (95%) cotransfected cells that were imaged, Vasa 463-661 relocalized to the plasma membrane in the presence of Oskar 139-240. ***, no obvious relocalization of mEGFP-Vasa 463-661 to the plasma membrane was detected, probably because the expression level of PH-mCherry-Oskar 139-240 was comparably low. In two additional replicates, we observed relocalization in 31 of 31 (100%) and in 22 of 23 (95%) cotransfected cells. The scale bar is 10 µm.

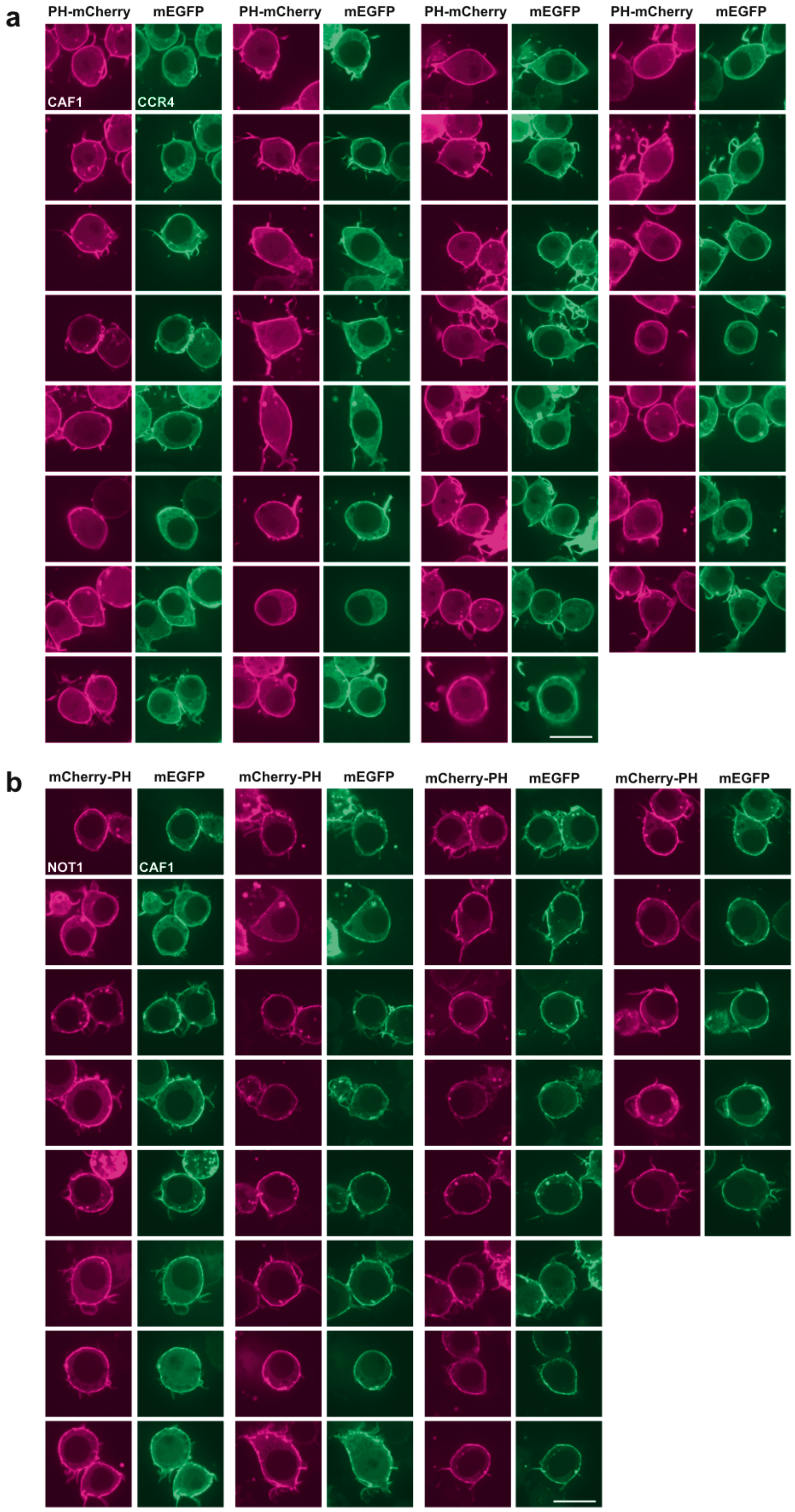


**Supplementary Fig. 3. Assessing the CCR4-CAF1 and CAF1-NOT1 interactions.** PH-mCherry, mCherry-PH, or mEGFP fusions to the proteins indicated were coexpressed in S2R+ cells and their localization was analyzed by microscopy. In 100% of the cells imaged an interaction between CCR4 and CAF1 (A; 31/31, 20/20, 45/45) or CAF1 and NOT1 (B; 29/29, 23/23, 30/30) was observed in three independent replicates each. The scale bar is 10 μm.

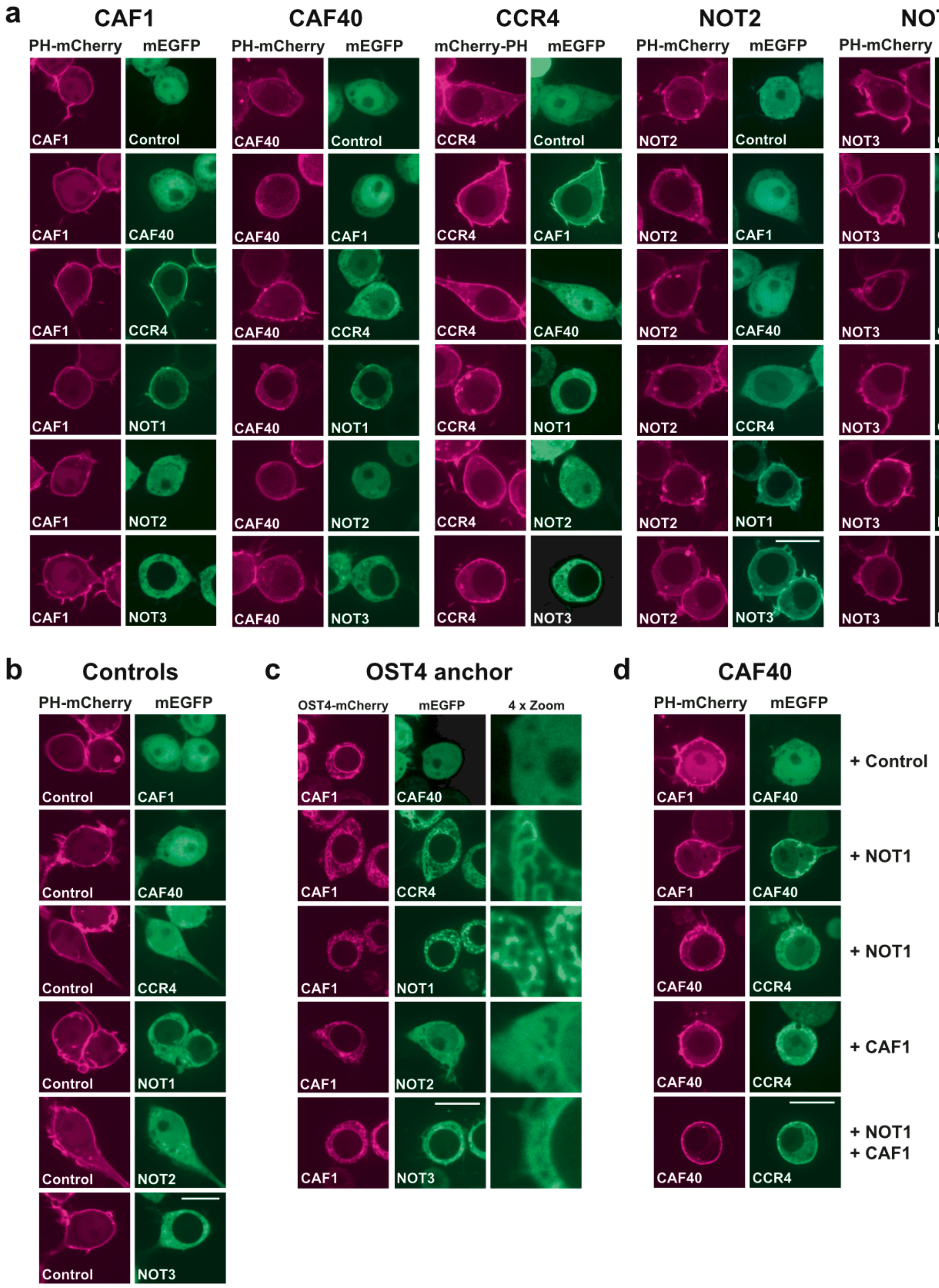


**Supplementary Fig. 4. Interactions between subunits of the CCR4-NOT complex.** PH-mCherry, OST4-mCherry, EGFP, or mEGFP fusions to the proteins indicated were coexpressed in S2R+ cells and their localization was analyzed by microscopy. **a** CAF1 recruited CCR4 and NOT1 to the plasma membrane but not CAF40, NOT2, or NOT3. CAF40 interacted with NOT1, and CCR4 interacted with CAF1. NOT2 bound to both NOT1 and NOT3. NOT3 interacted with both NOT1 and NOT2. **b** Localization of the core subunits of the CCR4-NOT complex fused to mEGFP in the presence of a PH-mCherry control plasmid. **c** PPIs between OST4 fusions of CAF1 and CCR4-NOT complex subunits. CAF1 recruited both CCR4 and NOT1 to the ER membrane, but not CAF40, NOT2, or NOT3. **d** CAF1 interacted with CAF40 upon NOT1 coexpression, and with CCR4 when both CAF1 and NOT1 were coexpressed. The scale bar is 10 µm.

a

| Subunit | Expression levels in S2R+ cells (RPKM) |
|---|---|
| Arp2 | 56 |
| Arp3 | 329 |
| Arpc1 | 263 |
| Arpc2 | 97 |
| Arpc3B | 24 |
| Arpc4 | 170 |
| Arpc5 | 49 |
| CAF1 | 28 |
| CAF40 | 74 |
| CCR4 | 16 |
| NOT1 | 47 |
| NOT2 | 48 |
| NOT3 | 87 |

No - low (0-10)
Moderate (11-25)
Moderately high (26-50)
High (51-100)
Very high (101-1000)

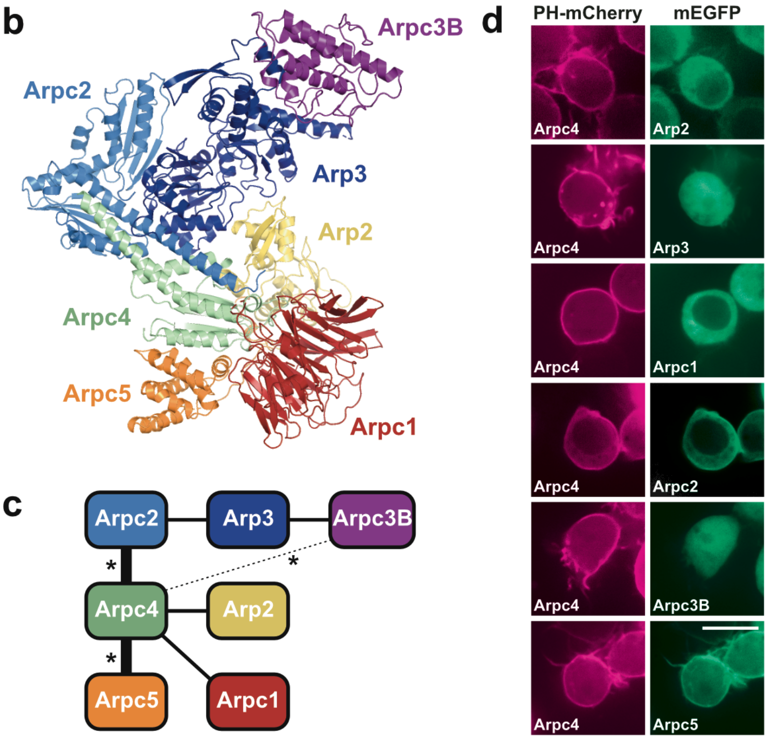


**Supplementary Fig. 5. Interactions between subunits of the Arp2/3 complex.**
**a** Expression levels of the subunits of the Arp2/3 and of the CCR4-NOT complex in S2R+ cells[1,2]. RPKM: reads per kilobase per million mapped bases. **b** Crystal structure of the bovine Arp2/3 complex[3]. **c** Scheme indicating the expected PPIs between the subunits of the Arp2/3 complex based on the crystal structure (straight connecting lines). Asterisks (*) indicate three interactions that were observed in pairwise Y2H tests with subunits of the human Arp2/3 complex[4]; of these three, the Arpc4 - Arpc3B interaction (dotted connecting line) is not expected based on the crystal structure. Connecting lines in bold represent PPIs that were observed in the ReLo assay. **d** PPI tests performed with the ReLo assay showing the results of the Arpc4 subunit, as an example. No interactions were observed when testing the subunits Arp2, Arp3, Arpc1, or Arpc3B (data not shown). PH-mCherry-Arpc4 and mEGFP fusions to the proteins indicated were coexpressed in S2R+ cells and their localization was analyzed by microscopy. Arpc4 interacted with Arpc2 and Arpc5. The scale bar is 10 μm.

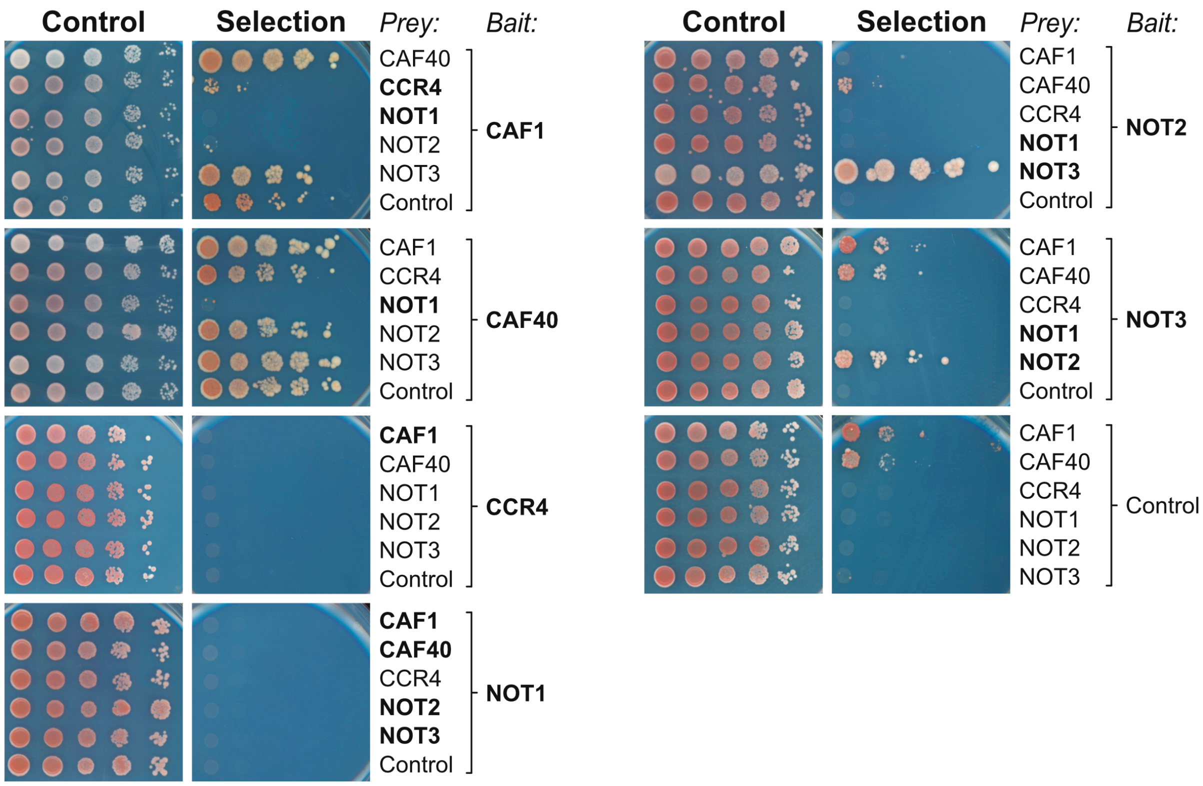


**Supplementary Fig. 6. Interaction tests between the subunits of the CCR4-NOT complex using Y2H.**
Split-ubiquitin yeast two-hybrid assays were performed with bait and prey constructs containing *Drosophila* proteins as indicated or no insertion (control plasmid). Five 10-fold dilutions of the cells were spotted and imaged after two (control plate) or six (selection plate) days of incubation. Selection medium lacked adenine and histidine. PPIs that were expected to be observed are indicated by protein names in bold letters.

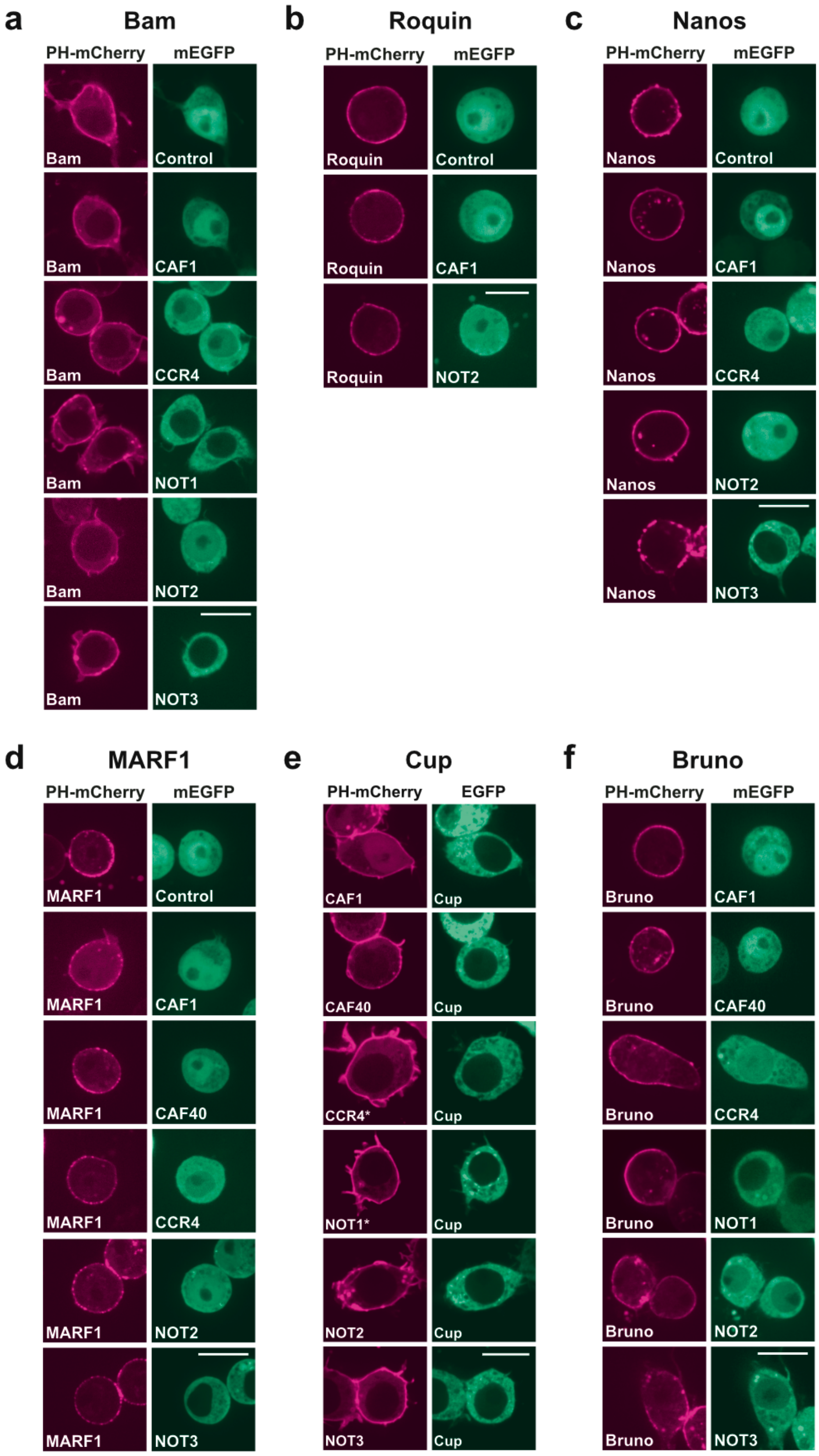


**Supplementary Fig. 7. Negative results obtained from PPI tests between the subunits of the CCR4-NOT complex and repressor proteins.**
PH-mCherry, mEGFP, or EGFP fusions to the proteins indicated were coexpressed in S2R+ cells and their localization was analyzed by microscopy. Shown are the negative results for the interaction tests between subunits of the CCR4-NOT complex and the proteins Bam (**a**), Roquin (**b**), Nanos, (**c**), MARF1 (**d**), Cup (**e**), and Bruno (**f**). *, CCR4 and NOT1 carried a C-terminal mCherry-PH fusion. The scale bar is 10 μm.

a

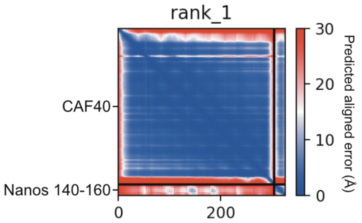


b

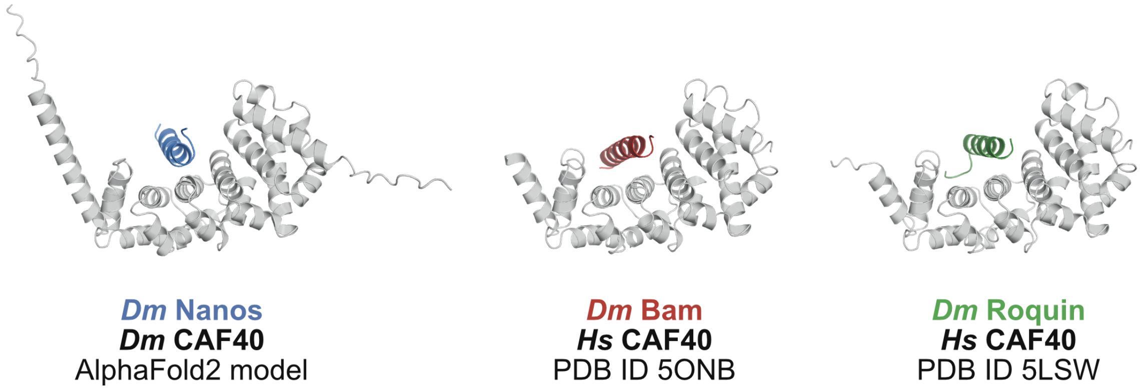


**Supplementary Fig. 8. Comparison of the predicted structure of Nanos-CAF40 to experimental structures of CAF40-peptide complexes.**

**a** Plot showing the predicted aligned error of the structural model obtained using AlphaFold2-Multimer version 3 and depicted in **Fig. 4h**. **b** *Drosophila* (*Dm*) or human (*Hs*) CAF40 (grey color) as indicated bound to *Drosophila* Nanos (blue color; obtained using AlphaFold2-Multimer version 3), *Drosophila* Bam (red color; [5]) or *Drosophila* Roquin (green color; [6])

**Supplementary Table 1. DNA constructs used in this study.**
All constructs are *Drosophila melanogaster* sequences, if not indicated otherwise. The specific protein isoforms (iso) used are indicated.

| **Vector** (insertion site) (code) | **Final DNA construct** | **DNA template information** | **Code** |
|---|---|---|---|
| **pAc5.1-EGFP** (EcoRV) (T5-MJ) | pAc5.1-EGFP-**Aubergine** iso A | *Drosophila* ovarian cDNA | F20-MJ |
| | pAc5.1-EGFP-**Aubergine 4R→K (**R11K/R13K/R15K/R17K) | Site directed mutagenesis of pAc5.1-EGFP-Aubergine | HK121 |
| | pAc5.1-EGFP-**Vasa** iso A | Reference: [7] | F15-MJ |
| | pAc5.1-EGFP-**Vasa 1-200** | pAc5.1-EGFP-Vasa | F17-MJ |
| | pAc5.1-EGFP-**Vasa 200-462** | pAc5.1-EGFP-Vasa | F18-MJ |
| | pAc5.1-EGFP-**Vasa 463-661** | pAc5.1-EGFP-Vasa | F19-MJ |
| | pAc5.1-EGFP-**Vasa 463-661 MUT** (F504E) | pAc5.1-EGFP-Vasa 463-661 | HK196 |
| | pAc5.1-EGFP-**Vasa closed** (E400Q) | Site directed mutagenesis of pAc5.1-EGFP-Vasa | F16-MJ |
| | pAc5.1-EGFP-**Vasa open** (K295N) | Site directed mutagenesis of pAc5.1-EGFP-Vasa | RR228 |
| | pAc5.1-EGFP-**Vasa F504E** | Site directed mutagenesis of pAc5.1-EGFP-Vasa | HK197 |
| | pAc5.1-EGFP-**Cup** iso B | pBSK-Cup-Flag (gift from Elmar Wahle) | F23-MJ |
| **pAc5.1-mEGFP** (EcoRV) (T6-MJ) | pAc5.1-mEGFP-**CAF1** iso A | pMTV5-Myc-CAF1 (gift from Elmar Wahle)[8] | HK50 |
| | pAc5.1-mEGFP-**CAF40** iso A | pET19-CAF40 (gift from Elmar Wahle) | HK51 |
| | pAc5.1-mEGFP-**CCR4** iso A | pMTV5-Myc-CCR4 (gift from Elmar Wahle) | HK52 |
| | pAc5.1-mEGFP-**NOT1** iso D | pSPL_Strep_NOT1_NOT2 (gift from Elmar Wahle) | HK53 |
| | pAc5.1-mEGFP-**NOT2** iso A | pSPL_Strep_NOT1_NOT2 (gift from Elmar Wahle) | HK54 |
| | pAc5.1-mEGFP-**NOT3** iso A | pFL-Flag-NOT3 (gift from Elmar Wahle) | HK55 |
| | pAc5.1-mEGFP-**human FRB** | pSF3-NBir-FKBP_CBir-FRB (gift from Julien Béthune)[9] | HK168 |
| | pAc5.1-mEGFP-**human MDM2 1-118** | Addgene clone 70413 (gift from Dominic Esposito) | HK181 |
| | pAc5.1-mEGFP-**Arp2** iso C | S2R+ cell cDNA | HK254 |
| | pAc5.1-mEGFP-**Arp3** iso A | *Drosophila* testis cDNA | HK265 |
| | pAc5.1-mEGFP-**Arpc1** iso A | *Drosophila* testis cDNA | HK268 |
| | pAc5.1-mEGFP-**Arpc2** iso A | S2R+ cell cDNA | HK255 |

| | | | |
|---|---|---|---|
| | pAc5.1-mEGFP-**Arpc3** iso C | S2R+ cell cDNA | HK256 |
| | pAc5.1-mEGFP-**Arpc5** iso A | S2R+ cell cDNA | HK258 |
| | pAc5.1-mEGFP-**CAF40 V186E** | pAc5.1-mEGFP-CAF40 | HK275 |
| **pAc5.1-mCherry** (EcoRV) (T7-MJ) (FspAI) (JM65) | pAc5.1-mCherry-**Oskar 139-606** | *oskar* cDNA | H2-MJ |
| | pAc5.1-mCherry-**Bruno** iso A | *bruno* cDNA | JM78 |
| | pAc5.1-mCherry-**Nanos** iso B | *nanos* cDNA | JM154 |
| **pAc5.1-λN-HA** (EcoRV) (T8-MJ) | pAc5.1-λN-HA-**CAF1** iso A | pMTV5-Myc-CAF1 (gift from Elmar Wahle)[8] | HK101 |
| | pAc5.1-λN-HA-**NOT1** iso D | pSPL_Strep_NOT1_NOT2 (gift from Elmar Wahle) | JM155 |
| | pAc5.1-λN-HA-**Cup** iso B | pBSK-Cup-Flag (gift from Elmar Wahle) | I14-MJ |
| **pAc5.1-PH-mEGFP** (FspAI) (JM50) | pAc5.1-PH-mEGFP-**Cup** iso B | pBSK-Cup-Flag (gift from Elmar Wahle) | JM51 |
| | pAc5.1-PH-mEGFP-**Bruno** iso A | *bruno* cDNA | JM165 |
| **pAc5.1-PH-mCherry** (FspAI) (HK49) | pAc5.1-PH-mCherry-**CAF1** iso A | pMTV5-Myc-CAF1 (gift from Elmar Wahle)[8] | HK96 |
| | pAc5.1-PH-mCherry-**CAF40** iso A | pET19-CAF40 (gift from Elmar Wahle) | HK97 |
| | pAc5.1-PH-mCherry-**NOT2** iso A | pSPL_Strep_NOT1_NOT2 (gift from Elmar Wahle) | HK99 |
| | pAc5.1-PH-mCherry-**NOT3** iso A | pFL-Flag-NOT3 (gift from Elmar Wahle) | HK100 |
| | pAc5.1-PH-mCherry-**human p53 1-50** | pcDNA3_1_3XHA_p53 WT (gift from Bernd Bukau) | HK180 |
| | pAc5.1-PH-mCherry-**human FKBP12** | pSF3-NBir-FKBP_CBir-FRB (gift from Julien Béthune)[9] | HK174 |
| | pAc5.1-PH-mCherry-**Oskar 139-606** iso A (Short Oskar) | pAc5.1-OST4-mCherry-Oskar 139-606 | HK195 |
| | pAc5.1-PH-mCherry-**Oskar 139-240** (eLOTUS domain) | pAc5.1-OST4-mCherry-Oskar 139-606 | HK73 |
| | pAc5.1-PH-mCherry-**Oskar 241-387** (DR) | pAc5.1-OST4-mCherry-Oskar 139-606 | HK74 |
| | pAc5.1-PH-mCherry-**Oskar 388-606** (OSK domain) | pAc5.1-OST4-mCherry-Oskar 139-606 | HK75 |
| | pAc5.1-PH-mCherry-**Oskar 139-240 MUT** (A162E/L228E) | Site directed mutagenesis of pAc5.1-PH-mCherry-Oskar 139-240 | HK202 |
| | pAc5.1-PH-mCherry-**Tudor** iso A | pAc5.1-EGFP-Tudor | HK130 |
| | pAc5.1-PH-mCherry-**Bam** iso A | *Drosophila* ovarian cDNA | HK62 |

| | | | |
|---|---|---|---|
| | pAc5.1-PH-mCherry-**Bam M24E** | Site directed mutagenesis of pAc5.1-PH-mCherry-Bam | HK88 |
| | pAc5.1-PH-mCherry-**Nanos** iso B | *nanos* cDNA | HK65 |
| | pAc5.1-PH-mCherry-**NanosΔ140-160** | pAc5.1 PH-mCherry-Nanos iso B | HK287 |
| | pAc5.1-PH-mCherry-**Roquin** iso A | *Drosophila* ovarian cDNA | HK63 |
| | pAc5.1-PH-mCherry-**MARF1** iso D | MARF1 cDNA[7] | HK56 |
| | pAc5.1-PH-mCherry-**Bruno** iso A | *bruno* cDNA | JM72 |
| | pAc5.1 PH-mCherry-**Arpc4** iso A | S2R+ cell cDNA | HK252 |
| **pAc5.1-mCherry-PH** (FspAI) (EB3) | pAc5.1-**CCR4** (iso A )-mCherry-PH | pMTV5-Myc-CCR4 (gift from Elmar Wahle) | EB7 |
| | pAc5.1-**NOT1** (iso D)-mCherry-PH | pSPL_Strep_NOT1_NOT2 (gift from Elmar Wahle) | EB5 |
| **pAc5.1-OST4-mCherry** (EcoRV) (XH26) | pAc5.1-OST4-mCherry-**CAF1** iso A | pMTV5-Myc-CAF1(gift from Elmar Wahle) | HK184 |
| | pAc5.1-OST4-mCherry-**Oskar 139-606** iso A | pAc5.1-mCherry-Oskar 139-606 | H3-MJ |
| | pAc5.1-OST4-mCherry-**Oskar 241-387** | pAc5.1-mCherry-Oskar 139-606 | HK193 |
| | pAc5.1-OST4-mCherry-**Oskar 388-606** | pAc5.1-mCherry-Oskar 139-606 | HK194 |
| **pDHB1-MJ** (Eco47III) (JK16) | pDHB1-MJ-**CAF1** iso A | pMTV5-Myc-CAF1 (gift from Elmar Wahle)[6] | JK19 |
| | pDHB1-MJ-**CAF40** iso A | pET19-CAF40 (gift from Elmar Wahle) | JK20 |
| | pDHB1-MJ-**CCR4** iso A | pMTV5-Myc-CCR4 (gift from Elmar Wahle) | JK21 |
| | pDHB1-MJ-**NOT1** iso D | pSPL_Strep_NOT1_NOT2 (gift from Elmar Wahle) | KM03 |
| | pDHB1-MJ-**NOT2** iso A | pSPL_Strep_NOT1_NOT2 (gift from Elmar Wahle) | JK32 |
| | pDHB1-MJ-**NOT3** iso A | pFL-Flag-NOT3 (gift from Elmar Wahle) | JK33 |
| **pPR3-N-MJ** (SmaI) (JK18) | pPR3-N-MJ-**CAF1** iso A | Reference: [10] | JK76 |
| | pPR3-N-MJ-**CAF40** iso A | Reference: [10] | JK77 |
| | pPR3-N-MJ-**CCR4** iso A | Reference: [10] | JK78 |
| | pPR3-N-MJ-**NOT1** iso D | pSPL_Strep_NOT1_NOT2 (gift from Elmar Wahle) | HK225 |
| | pPR3-N-MJ-**NOT2** iso A | Reference: [10] | JK88 |
| | pPR3-N-MJ-**NOT3** iso A | Reference: [10] | JK89 |

**Sequence of the PH domain of *Rattus norvegicus* PLCδ:**
HGLQDDPDLQALLKGSQLLKVKSSSWRRERFYKLQEDCKTIWQESRKVMRSPESQ
LFSIEDIQEVRMGHRTEGLEKFARDIPEDRCFSIVFKDQRNTLDLIAPSPADAQHWV
QGLRKIIHHSGSMDQRQK

**Sequence of the *Saccharomyces cerevisiae* OST4 miniprotein:**
MISDEQLNSLAITFGIVMMTLIVIYHAVDSTMSPKN

**Linker sequences between fluorescence protein and protein of interest:**
PSLNSATC (all ReLo vectors with both N-terminal tag and containing the PH sequence)
PSLNSAD (all other ReLo vectors with N-terminal tag)
ASSGGTN (all ReLo vectors with C-terminal tag)

### SUPPLEMENTARY METHODS

#### Yeast two-hybrid

Split-ubiquitin yeast-two hybrid assays were performed as described previously[11]. 500 ng each of bait pPR3-N and prey pDHB1 vectors were cotransformed into competent NMY51 yeast cells, which were then plated onto SDC agar lacking leucine and tryptophan and incubated at 30°C for two days. To perform the spotting assay, three to five colonies were picked, resuspended in water, and the cell suspension was then diluted to an $OD_{600}$ of 3 and to four more consecutive 1:10 dilutions. 5 µl of each dilution was spotted on SDC agar plates either lacking leucine and tryptophan (control plates) or lacking leucine, tryptophan, adenine, and histidine (selection plate). The plates were then incubated for two (control plate) or six (selection plate) days at 30°C and images were taken.

#### Structural prediction

For the prediction of the structure of the Nanos-CAF40 complex the ColabFold v1.5.2 web interface[12] was used with standard settings except for the model_type, which was switched from "auto" to "alphaFold2_multimer_v3". Structures were visualized using Pymol.
